# Modern-Day SIV Viral Diversity Generated by Extensive Recombination and Cross-Species Transmission

**DOI:** 10.1101/101725

**Authors:** Sidney M. Bell, Trevor Bedford

**Affiliations:** Vaccine and Infectious Disease Division, Fred Hutchinson Cancer Research Center, Seattle, Washington, United States of America; Molecular and Cellular Biology Program, University of Washington, Seattle, Washington, United States of America

## Abstract

Cross-species transmission (CST) has led to many devastating epidemics, but is still a poorly understood phenomenon. HIV-1 and HIV-2 (human immunodeficiency virus 1 and 2), which have collectively caused over 35 million deaths, are the result of multiple CSTs from chimpanzees, gorillas, and sooty mangabeys. While the immediate history of HIV is known, there are over 45 lentiviruses that infect specific species of primates, and patterns of host switching are not well characterized. We thus took a phylogenetic approach to better understand the natural history of SIV recombination and CST. We modeled host species as a discrete character trait on the viral phylogeny and inferred historical host switches and the pairwise transmission rates between each pair of 24 primate hosts. We identify 14 novel, well-supported, ancient cross-species transmission events. We also find that lentiviral lineages vary widely in their ability to infect new host species: SIV*col* (from colobus monkeys) is evolutionarily isolated, while SIV*agm*s (from African green monkeys) frequently move between host subspecies. We also examine the origins of SIV*cpz* (the predecessor of HIV-1) in greater detail than previous studies, and find that there are still large portions of the genome with unknown origins. Observed patterns of CST are likely driven by a combination of ecological circumstance and innate immune factors.

## Introduction

As demonstrated by the recent epidemics of EBOV and MERS, and by the global HIV pandemic, viral cross-species transmissions (CST) can be devastating [1,2]. As such, understanding the propensity and ability of viral pathogens to cross the species barrier is of vital public health importance [3]. Of particular interest are transmissions that not only “spillover” into a single individual of a new host species, but that result in a virus actually establishing a sustained chain of transmission and becoming endemic in the new host population (“host switching”).

HIV is the product of not just one successful host switch, but a long chain of host switch events [4,5]. There are two human immunodeficiency viruses, HIV-2 and HIV-1. HIV-2 arose from multiple cross-species transmissions of SIV*smm* (simian immunodeficiency virus, *sooty mangabey*) from sooty mangabeys to humans [6–8]. HIV-1 is the result of four independent cross-species transmissions from chimpanzees and gorillas. Specifically, SIV*cpz* was transmitted directly from chimpanzees to humans twice; one of these transmissions generated HIV-1 group M, which is the primary cause of the human pandemic [9]. SIV*cpz* was also transmitted once to gorillas [10], generating SIV*gor*, which was in turn transmitted twice to humans [11].

Looking further back, SIV*cpz* itself was also generated by lentiviral host switching and recombination. Based on the SIV sequences available at the time, early studies identified SIV*mon/-mus/-gsn* (which infect mona, mustached, and greater spot-nosed monkeys, respectively) and SIV*rcm* (which infects red-capped mangabeys) as probable donors [12]. Functional analysis of accessory genes from these putative parental lineages indicate that the specific donors and genomic locations of these recombination event(s) were crucial for enabling what became SIV*cpz* to cross the high species barrier and establish an endemic lineage in hominids [13].

The complex evolutionary history resulting in HIV illustrates the importance of natural history to modern day viral diversity, and although the history leading to HIV is well detailed, broader questions regarding cross-species transmission of primate lentiviruses remain [14]. With over 45 known extant primate lentiviruses, each of which is endemic to a specific host species [4,5,15], we sought to characterize the history of viral transmission between these species to the degree possible given the precision allowed from a limited sample of modern-day viruses.

Here, we utilized phylogenetic inference to reconstruct the evolutionary history of primate lentiviral recombination and cross-species transmission. We assembled datasets from publicly available lentiviral genome sequences and conducted discrete trait analyses to infer rates of transmission between primate hosts. We find evidence for extensive interlineage recombination and identify many novel host switches that occurred during the evolutionary history of lentiviruses. We also find that specific lentiviral lineages exhibit a broad range of abilities to cross the species barrier. Finally, we also examined the origins of each region of the SIV*cpz* genome in greater detail than previous studies to yield a more nuanced understanding of its origins.

## Results

### There have been at least 13 interlineage recombination events that confuse the lentiviral phylogeny

In order to reconstruct the lentiviral phylogeny, we first had to address the issue of recombination, which is frequent among lentiviruses [16]. In the context of studying cross-species transmission, this is both a challenge and a valuable tool. Evidence of recombination between viral lineages endemic to different hosts is also evidence that at one point in time, viruses from those two lineages were in the same animal (i.e., a cross-species transmission event must have occurred in order to generate the observed recombinant virus). However, this process also results in portions of the viral genome having independent evolutionary—and phylogenetic—histories.

To address the reticulate evolutionary history of SIVs, we set out to identify the extent and nature of recombination between lentiviral lineages. Extensive sequence divergence between lineages masks site-based methods for linkage estimation (Fig. S1). However, topology-based measures of recombination allow for “borrowing” of information across nearby sites, and are effective for this dataset. We thus utilized a phylogenetic model to group segments of shared ancestry separated by recombination breakpoints instantiated in the HyPhy package GARD [17]. For this analysis, we used a version of the SIV compendium alignment from the Los Alamos National Lab (LANL), modified slightly to reduce the overrepresentation of HIV sequences (N=64, see Methods) [18]. Importantly, because each virus lineage has only a few sequences present in this alignment, these inferences refer to *inter-*lineage recombination, and not the rampant *intra-*lineage recombination common among lentiviruses.

GARD identified 13 locations along the genome that had strong evidence of inter-lineage recombination (Fig. 1). Here, evidence for a particular model is assessed via Akaike Information Criterion (AIC) [19] and differences in AIC between models indicate log probabilities, so that a delta-AIC of 10 between two models would indicate that one model is e^10/2^ = ~148 as likely as the other. In our case, delta-AIC values ranged from 154 to 436 for each included breakpoint, indicating that these breakpoints are strongly supported by the underlying tree likelihoods. The 14 resulting segments ranged in length from 351 to 2316 bases; in order to build reliable phylogenies, we omitted two of the less supported breakpoints from downstream analyses, yielding 12 segments ranging in length from 606 – 2316 bases. We found no evidence to suggest linkage between non-neighboring segments (Fig. S2). While it has been previously appreciated that several lineages of SIV are recombinant products [12,20,21], the 13 breakpoints identified here provide evidence that there have been at least 13 inter-lineage recombination events during the evolution of SIVs. Identifying these recombination breakpoints allowed us to construct a putatively valid phylogeny for each segment of the genome that shares an internally cohesive evolutionary history.

**Figure 1:**
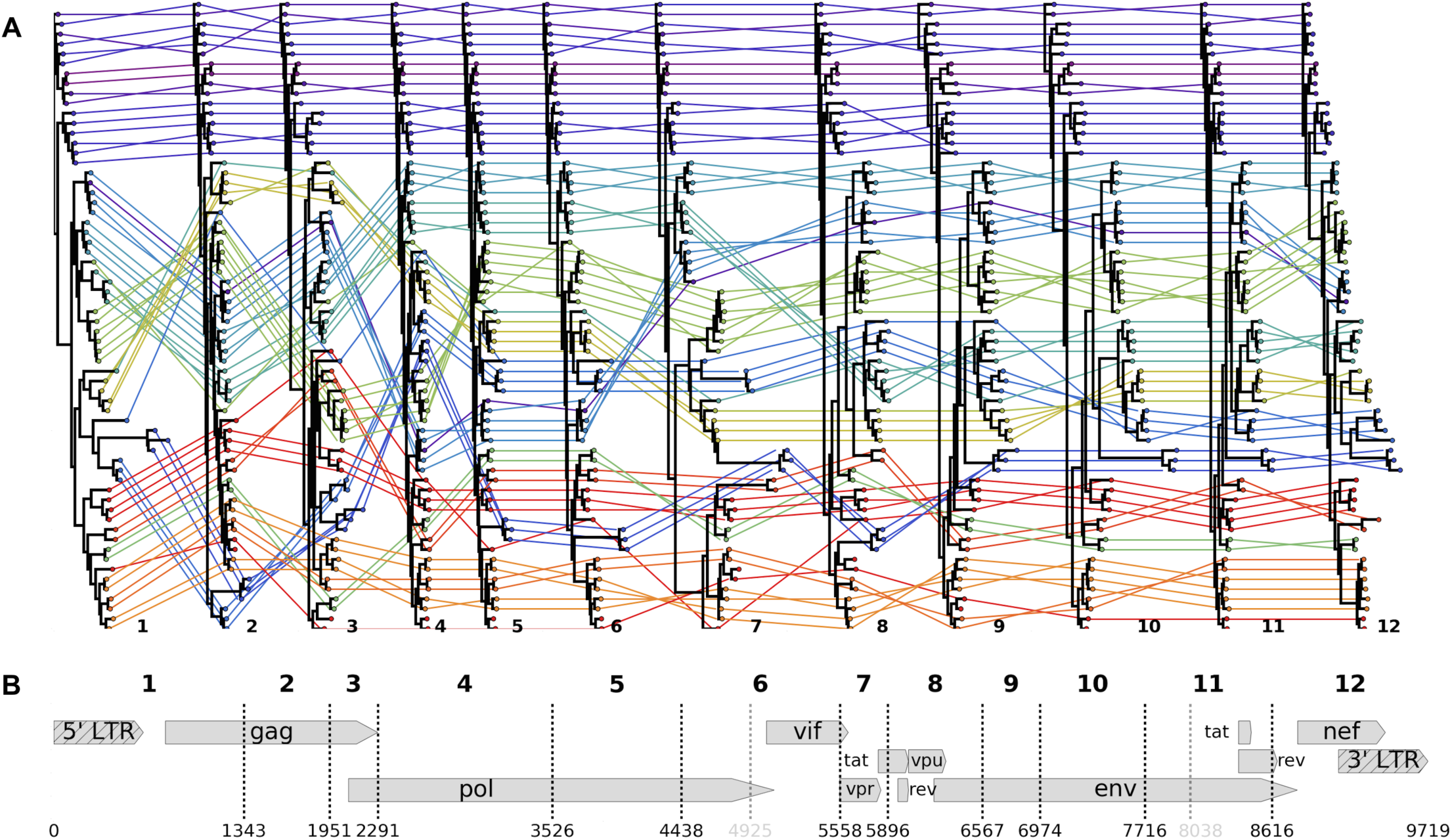
There have been at least 13 interlineage recombination events among SIVs. The SIV LANL compendium, slightly modified to reduce overrepresentation of HIV, was analyzed with GARD to identify the 13 recombination breakpoints across the genome (dashed lines in **B**; numbering according to the accepted HXB2 reference genome-­‐-­‐accession K03455, illustrated). Two of these breakpoints were omitted from further analyses because they created extremely short fragments (< 500 bases; gray dashes in **B**). For each of the 11 remaining breakpoints used in further analyses, we split the compendium alignment along these breakpoints and built a maximum likelihood tree, displayed in **A.** Each viral sequence is color-­‐coded by host species, and its phylogenetic position is traced between trees. Heuristically, straight, horizontal colored lines indicate congruent topological positions between trees (likely not a recombinant sequence); criss-­‐crossing colored lines indicate incongruent topological positions between trees (likely a recombinant sequence).

### Most primate lentiviruses were acquired by cross-species transmissions

We then looked for phylogenetic evidence of cross-species transmission in the tree topologies of each of the 12 genomic segments. For this and all further analyses, we constructed a dataset from all publicly available primate lentivirus sequences, which we curated and subsampled by host and virus lineage to ensure an equitable distribution of data (see Methods). This primary dataset consists of virus sequences from the 24 primate hosts with sufficient data available (5 – 25 sequences per viral lineage, N=423, Fig. S3). Alignments used the fixed compendium alignment as a template (see Methods).

In phylogenetic trees of viral sequences, cross-species transmission appears as a mismatch between the host of a virus and the host of that virus’s ancestor. To identify this pattern and estimate how frequently each pair of hosts has exchanged lentiviruses, we used the established methods for modeling evolution of discrete traits, as implemented in BEAST [22,23]. In our case, the host of each viral sample was modeled as a discrete trait. This is analogous to treating the “host state” of each viral sample as an extra column in an alignment, and inferring the rate of transition between all pairs of host states along with inferring ancestral host states across the phylogeny. This approach is similar to common phylogeographic approaches that model movement of viruses across discrete spatial regions [24] and has previously been applied to modeling discrete host state in the case of rabies virus [22]. Here, we took a fully Bayesian approach and sought the posterior distribution across phylogenetic trees, host transition rates and ancestral host states. We integrate across model parameters using Markov chain Monte Carlo (MCMC). The resulting model provides phylogenetic trees for each segment annotated with ancestral host states alongside inferred transition rates.

Figure 2 shows reconstructed phylogenies for 3 segments along with inferred ancestral host states. Trees are color coded by known host state at the tips, and inferred host state at internal nodes/branches; color saturation indicates the level of certainty for each ancestral host assignment. A visual example of how the model identifies cross-species transmissions can be seen in the SIV*mon*/SIV*tal* clade, which infect mona- and talapoin monkeys, respectively (starred in Fig. 2A-C). Due to the phylogenetic placement of the SIV*mon* tips, the internal node at the base of this clade is red, indicating that the host of the ancestral virus was most likely a mona monkey. This contrasts with the host state of the samples isolated from talapoin monkeys (tips in green). These changes in the host state across the tree are what inform our estimates of the rate of transmission between host pairs. In total, the support for each possible transmission is derived from both A) whether the transmission is supported across the posterior distribution of phylogenies for a particular segment, and B) whether this is true for multiple genomic segments.

**Figure 2:**
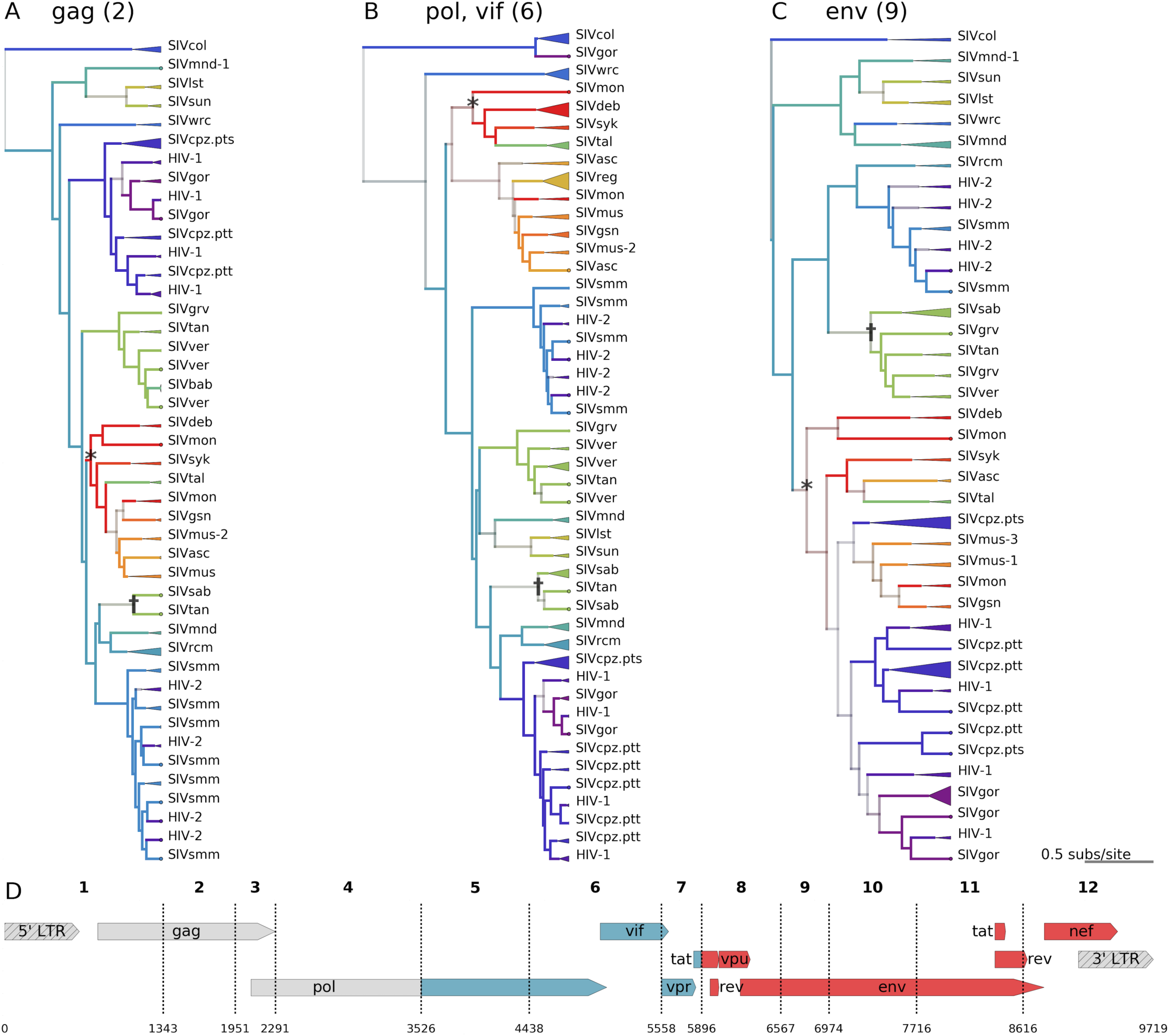
Cross-species transmissions are inferred from tree topologies; SIVcpz has mosaic origins. **A, B, C** -­‐ Bayesian maximum clade credibility (mcc) trees are displayed for segments 2 (*gag* − **A**), 6 (*int* and *vif* -­‐**B**), and 9 (*env* −**C**) of the main dataset (N=423). Tips are color coded by known host species; internal nodes and branches are colored by inferred host species, with saturation indicating the confidence of these assignments. Monophyletic clades of viruses from the same lineage are collapsed, with the triangle width proportional to the number of represented sequences. An example of likely cross-­‐species transmission is starred in each tree, where the host state at the internal node (red / mona monkeys) is incongruent with the descendent tips’ known host state (green / talapoin monkeys), providing evidence for a transmission from mona monkeys to talapoin monkeys. Another example of cross-­‐species transmission of a recombinant virus among African green monkeys is marked with a dagger. **D** − The genome map of SIVcpz, with breakpoints used for the discrete trait analysis, is color coded and labeled by the most likely ancestral host for each segment of the genome.

Notably, the tree topologies are substantially different between segments, which emphasizes both the extent of recombination and the different evolutionary forces that have shaped the phylogenies of individual portions of the genome. In all segments’ trees, we also see frequent changes in the host state between internal nodes (illustrated as changes in color going up the tree), suggestive of frequent ancient cross-species transmissions. On average, primate lentiviruses switch hosts once every 6.25 substitutions per site per lineage across the SIV phylogeny.

The cross-species transmission events inferred by the model are illustrated in Fig. 3, with raw rates and Bayes factors (BF) in Fig. S4. As shown, the model correctly infers the pairs of hosts with previously identified CST events [6,9,11,12,20,21,25,26]. Importantly, we also identify 14 novel cross-species transmission events with strong statistical support (cutoff of BF >= 10.0). Each of these transmissions is clearly and robustly supported by the tree topologies (all 12 trees are illustrated in Fig. S5).

**Figure 3:**
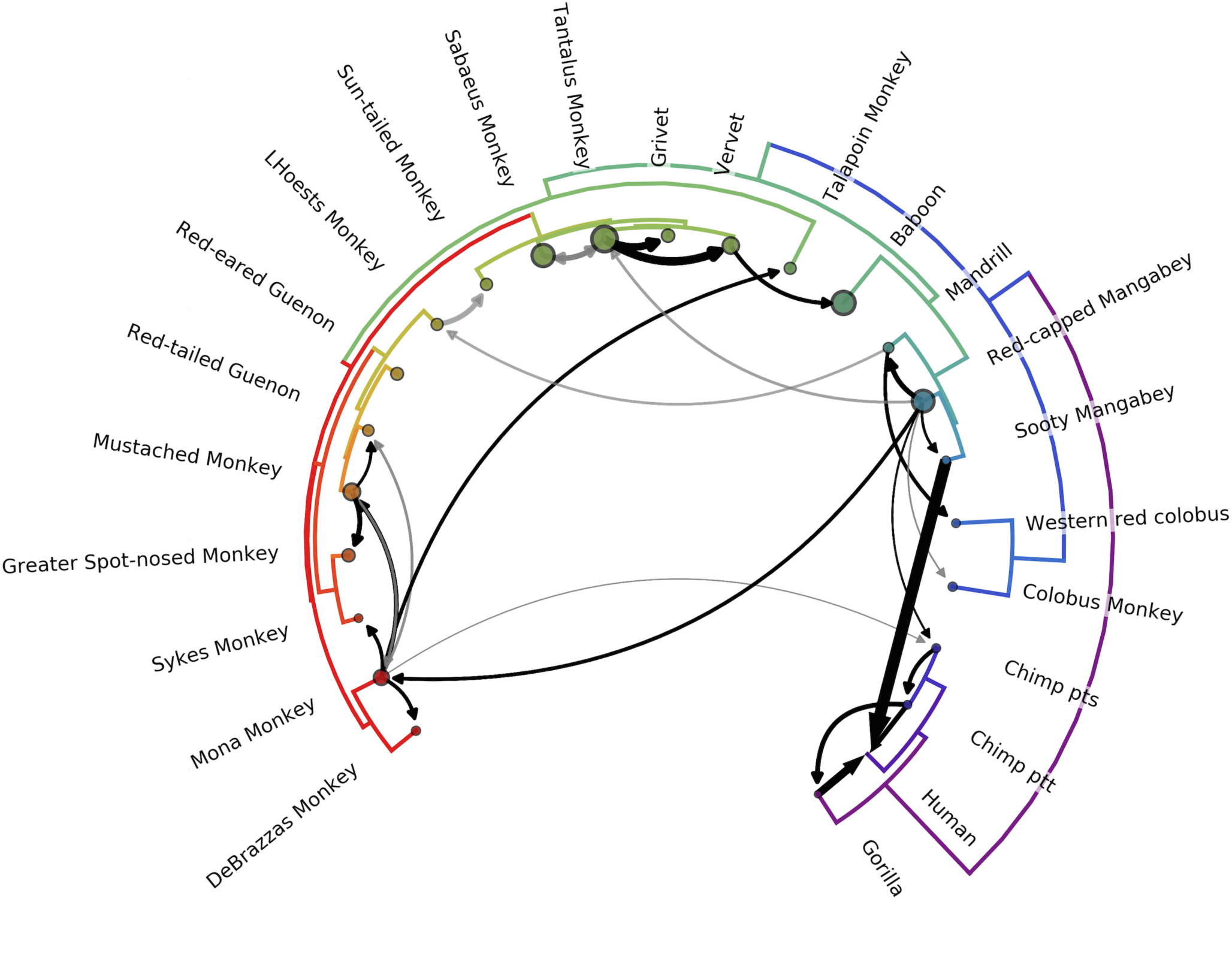
Most lentiviruses are the product of ancient cross-species transmissions. The phylogeny of the host species’ mitochondrial genomes forms the outer circle. Arrows represent transmission events inferred by the model with Bayes’ factor (BF) >= 3.0; black arrows have BF >= 10, with opacity of gray arrows scaled for BF between 3.0 and 10.0. Width of the arrow indicates the rate of transmission (actual rates = rates * indicators). Circle sizes represent network centrality scores for each host. Transmissions from chimps to humans; chimps to gorillas; gorillas to humans; sooty mangabeys to humans; sabaeus to tantalus; and vervets to baboons have been previously documented. To our knowledge, all other transmissions illustrated are novel identifications.

To control for sampling effects, we repeated the analysis with a supplemental dataset built with fewer hosts, and more sequences per host (15 host species, subsampled to 16-40 sequences per viral lineage, N=510), and see consistent results. As illustrated in Figs. S6-8, we find qualitatively similar results. When directly comparing the average indicator values, host state transition rates, and BF values between analyses, the results from the main and supplemental datasets are strongly correlated, indicating robust quantitative agreement between the two analyses (Fig. S9). Taken together, these results represent a far more extensive pattern of CST among primate lentiviruses than previously described, with all but a few lineages showing robust evidence of host switching [4,5,14].

However, the distribution of these host switches is not uniform; when we assess the network centrality of each virus we find a broad range (Figs. 3, S6, as node size), indicating some hosts act as sources in the SIV transmission network and other hosts act as sinks. From this, we infer that some viruses have either had greater opportunity or have a greater ability to cross the species barrier than others. In particular, the SIVs from four of the subspecies of African green monkeys (SIV*sab*, SIV*tan*, SIV*ver*, SIV*pyg*; collectively, SIV*agm*) appear to exchange viruses with other host species frequently (Fig. 3, 12’ o-clock). An example of SIV*agm* CST events can be seen in the tree topologies from *gag, prot, reverse transcriptase (RT)*, and *vif* (Figs. 2 and S5, segments 2, 3, 4, & 6). Here, SIV*tan* isolates reported by Jin *et al*. (denoted with ✝ in Fig. 2) clearly cluster with SIV*sab*, in a distant part of the phylogeny from the rest of the SIV*agm* viruses (including the majority of SIV*tan* isolates) [20]. For all other segments, however, the SIV*agm*s cluster together. We thus concur with the conclusion of Jin *et al.* that these samples represent a recent spillover of SIV*sab* from sabaeus monkeys to tantalus monkeys, and the model appropriately identifies transmission.

Although SIV*agm*s appear to be particularly prone to CST among African green monkeys, nearly every primate clade has at least one inbound, robustly supported viral transmission from another clade. We thus conclude that the majority of lentiviruses have arisen from a process of host switching, followed by a combination of intraclade host switches and host-virus coevolution.

### SIV*col* appears to be evolutionarily isolated

Previous studies of the lentiviral phylogeny have noted that SIV*col* is typically the outgroup to other viral lineages, and have hypothesized that this may implicate SIV*col* as the “original” primate len-tivirus [27]. We find this hypothesis plausible, but the evidence remains inconclusive. For the majority of genomic segments, we also observe SIV*col* as the clear outgroup (Figs. 2 and S5). In contrast, for portions of *gag/pol* (segments 3 and 4) and some of the accessory genes (segment 7), we find that there is not a clear outgroup. For these segments, many other lineages of SIV are just as closely related to SIV*col* as they are to each other. However, with the occasional exception of single heterologous taxa with poorly supported placement, SIV*col* remains a monophyletic clade (N=16), and does not intercalate within the genetic diversity of any other lineage in our dataset.

Based on these collective tree topologies, our model does not identify strong evidence for any specific transmissions out of colobus monkeys, and identifies only a single, weakly supported inbound transmission (likely noise in the model caused by the fact that red-capped mangabeys are the marginally supported root host state; see below). This is consistent with previous findings that the colobinae have a unique variant of the *APOBEC3G* gene, which is known to restrict lentiviral infection and speculated to be a barrier to cross-species transmission [27]. These observations generally support the idea of SIV*col* as having maintained a specific relationship with its host over evolutionary time. However, we find inconclusive evidence for the hypothesis that SIV*col* is the most ancient or “source” lineage of primate lentivirus; if this hypothesis were true, then these lentiviral transmissions out of the colobinae must have been followed by extensive recombination that obscures the relationships between SIV*col* and its descendants.

### SIV*cpz*, the precursor to HIV-1, has a mosaic origin with unknown segments

Unlike SIV*col*, SIV*cpz* appears to be the product of multiple CSTs and recombination events. SIV*cpz* actually encompasses two viral lineages: SIV*cpzPtt* infects chimpanzees of the subspecies *Pantroglodytes troglodytes*, and SIV*cpzPts* infects chimpanzees of the subspecies *Pan troglodytes schweinfurthii* [28]. There are two additional subspecies of chimpanzees that have not been found harbor an SIV despite extensive surveys, suggesting that SIV*cpzPtt* was acquired after chimpanzee subspeciation [26]. Both previous work [26,28] and our own results support the hypothesis that SIV*cpz* was later transmitted from one chimpanzee subspecies to the other, and SIV*cpzPtt* is the only SIV*cpz* lineage that has crossed into humans. Given the shared ancestry of the two lineages of SIV*cpz*, we use “SIV*cpz*” to refer specifically to SIV*cpzPtt*.

Based on the lentiviral sequences available in 2003, Bailes et al [12] conclude that the SIV*cpz* genome is a recombinant of just two parental lineages. SIV*rcm* (which infects red-capped mangabeys) was identified as the 5’ donor, and an SIV from the SIV*mon/-mus/-gsn* clade (which infect primates in the *Cercopithecus* genus) was identified as the 3’ donor. Since the time of this previous investigation many new lentiviruses have been discovered and sequenced. In incorporating these new data, we find clear evidence that the previous two-donor hypothesis may be incomplete.

The tree topologies from *env* in the 3’ end of the genome (segments 8-11) support the previous hypothesis [12] that this region came from a virus in the SIV*mon/-mus/-gsn* clade. These viruses form a clear sister clade to SIV*cpz* with high posterior support (Fig. 2C,D). We find strong evidence for transmissions from mona monkeys (SIV*mon*) to mustached monkeys (SIV*mus*), and from mustached monkeys to greater spot-nosed monkeys (SIV*gsn*) (see discussion of potential coevolution below). We also find more evidence in support of a transmission from mona monkeys to chimpanzees than from the other two potential donors, but more sampling is required to firmly resolve which of these viruses was the original donor of the 3’ end of SIV*cpz*.

We find phylogenetic evidence to support the previous hypothesis [12,13] that the *integrase* and *vif* genes of SIV*cpz* (segments 4-6) originated from SIV*rcm*; however, we find *equally strong evidence* to support the competing hypothesis that *pol* came from SIV*mnd-2*, which infects mandrils (Fig. 2B,D). In these portions of the genome, SIV*mnd-2* and SIV*rcm* together form a clear sister clade to SIV*cpz*. The *vpr* gene, in segment 7, is also closely related to both SIV*rcm* and SIV*mnd-2*, but this sister clade also contains SIV*smm* from sooty mangabeys.

Interestingly, we do *not* find evidence to support either SIV*rcm/mnd-2* or SIV*mon/mus/gsn* as the donor for the 5’ most end of the genome (segments 1-5), including the *5’ LTR, gag*, and *RT* genes. This is also true for the *3’ LTR* (segment 12). SIV*cpz* lacks a clear sister clade or ancestor in this region, and SIV*rcm* groups in a distant clade; we therefore find no evidence to suggest that an ancestor of an extant SIV*rcm* was the parental lineage of SIV*cpz* in the 5’ most end of the viral genome as previously believed (Fig. 2A,D). This may support the possibility of a third parental lineage, or a number of other plausible scenarios (discussed below).

## Discussion

### Limitations and strengths of the model

#### Additional sampling is required to fully resolve the history of CST among lentiviruses

In addition to the 14 strongly supported novel transmissions (BF >= 10) described above, we also find substantial evidence for an additional 8 possible novel transitions, but with lower support (BF >= 3) (Fig. 3, S4). These transmissions are more difficult to assess, because many of them are inferred on the basis of just a few “outlier” tips of the tree that group apart from the majority of viral samples from the same lineage. In each case, the tips’ phylogenetic position is strongly supported, and the primary literature associated with the collection of each of these “outlier” samples clearly specifies the host metadata. However, due to the limited number of lentiviral sequences available for some hosts, we are unable to control for sampling effects for some of these lower-certainty transmissions. We report them here because it is unclear whether these outliers are the result of unidentified separate endemic lineages, one-time spillovers from other hosts, or species misidentification during sample collection. It is also important to note that while some of these less-supported transmissions are potentially sampling artifacts, many of them may be real, and may be less supported simply because they lack the requisite available data for some genome segments. Ultimately, far more extensive sampling of primate lentivi-ruses is required in order to resolve these instances.

#### Most lentiviruses were originally acquired by CST and have since coevolved with their hosts

Some of these “noisier” transmission inferences, particularly within the same primate clade, may be the result of coevolution, ie. lineage tracking of viral lineages alongside host speciation. Within the model, viral jumps into the *common ancestor* of two extant primate species appear as a jump into *one* of the extant species, with a secondary jump between the two descendants. For example, the model infers a jump from mona monkeys into mustached monkeys, with a secondary jump from mustached monkeys into their sister species, red-tailed guenons (Fig. 3). Comparing the virus and host phylogenies, we observe that this host tree bifurcation between mustached monkeys and red-tailed guenons is mirrored in the virus tree bifurcation between SIV*mus* and SIV*rtg* for most segments of the viral ge-nome (Figs. 2, 3). This heuristically suggests that the true natural history may be an ancient viral transmission from mona monkeys into the common ancestor of mustached monkeys and red-tailed guenons, followed by host/virus coevolution during primate speciation to yield SIV*mus* and SIV*rtg*.

The possibility of virus/host coevolution means that while we also observe extensive host switching *between* primate clades, many of the observed jumps *within* a primate clade may be the result of host-virus coevolution. However, we also note that the species barrier is likely lower between closely related primates, making it challenging to rigorously disentangle coevolution vs. true host switches within a primate clade [29].

#### The model propagates and accounts for residual uncertainty from the recombination analyses

The deep phylogenies and extensive sequence divergence between SIV lineages makes any assessment of recombination imperfect. Our analysis aims to 1) place a lower bound on the number of interlineage recombination events that must have occurred to explain the observed extant viruses, and 2) use this understanding to construct a model of CST among these viruses. As the most well developed package currently available for topology-based recombination analysis, GARD was an appropriate choice to identify in broad strokes the extent and nature of recombination among SIVs. Some (though not all) of the remaining uncertainty as to the exact location of breakpoints is represented within the posterior distribution of trees for each segment, and is thus propagated and accounted for in the discrete traits model (inferences of CST). Most other studies of SIV evolutionary history simply split the phylogeny along gene boundaries or ignore recombination (e.g., [5,12,29]). Thus, while there is still some remaining uncertainty in our recombination analyses, these results still represent a major step forward in attempting to systematically assess recombination among all extant SIV lineages and to incorporate it into the phylogenetic reconstruction.

### Cross-species transmission is driven by exposure and constrained by host genetic distance

Paleovirology and, more recently, statistical models of lentiviral evolution estimate that primate lentiviruses share a common ancestor approximately 5-10 million years ago [27,30,31]. This, along withthe putative viral coevolution during primate speciation, suggests that many of these transmissions were ancient, and have been acted on by selection for millions of years. Thus, given that the observed transmissions almost exclusively represent evolutionarily successful host switches, it is remarkable that lentiviruses have been able to repeatedly adapt to so many new host species. In the context of this vast evolutionary timescale, however, we conclude that while lentiviruses have a far more extensive history of host switching than previously understood, these events remain relatively rare overall.

Looking more closely at the distribution of host switching among lineages, it is clear that some viruses have crossed the species barrier with far greater frequency than others. This is likely governed by both ecological and biological factors. Ecologically, frequency and form of exposure are likely key determinants of transmission, but these relationships can be difficult to describe statistically [3]. For example, many primates are chronically exposed to many exogenous lentiviruses through predation [32]. Using log body mass ratios [33] as a proxy for predation, we do not see a statistically significant association between body mass ratio and non-zero transmission rate (Fig. 4A, blue; p=0.678, coef. 95% CI (-0.311, 0.202)). We believe the lack of signal is likely due to the imperfect proxy, although it is also possible that predator-prey relationships do not strongly structure the CST network.

**Figure 4:**
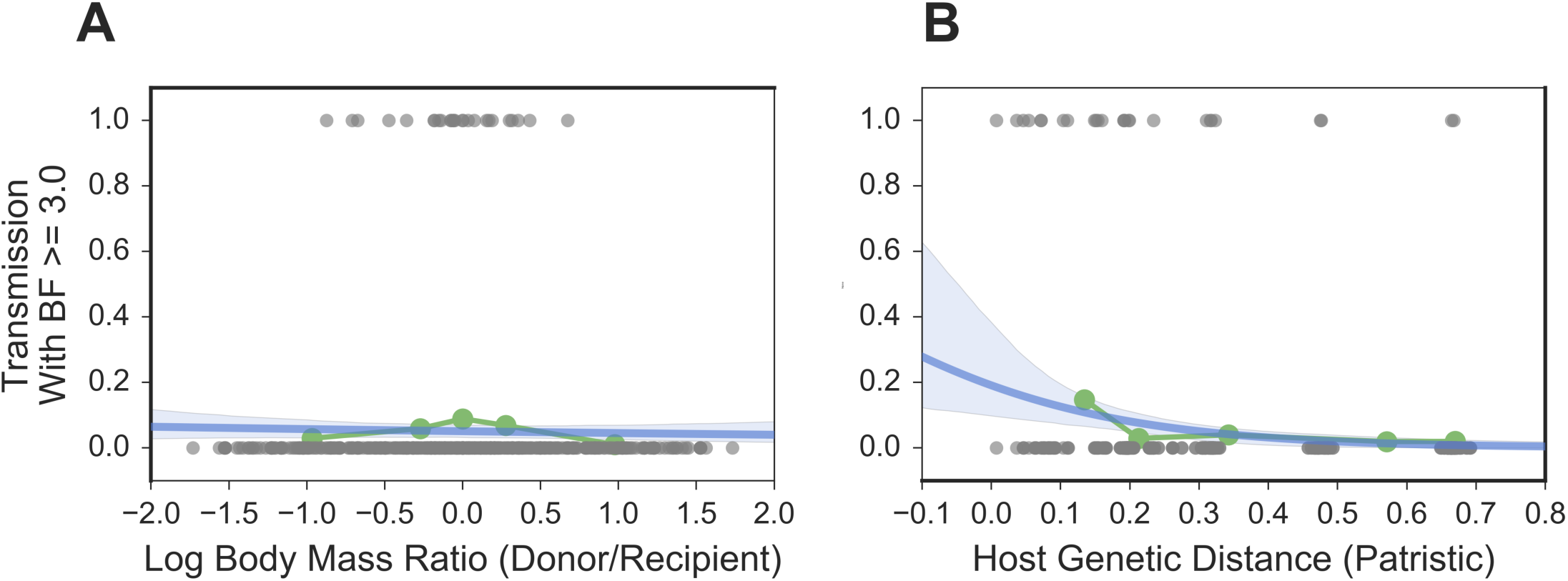
Cross-species transmission is driven by exposure and constrained by host genetic distance. For each pair of host species, we (**A**) calculated the log ratio of their average body masses and (**B**) found the patristic genetic distance between them (from a maximum-­‐likelihood tree of mtDNA). To investigate the association of these predictors with cross-­‐species transmission, we treated transmission as a binary variable: 0 if the Bayes factor for the transmission (as inferred by the discrete traits model) was < 3.0, and 1 for a Bayes factor >= 3.0. Each plot shows raw predictor data in gray; the quintiles of the predictor data in green; and the logistic regression and 95% CI in blue.

Biologically, increasing host genetic distance has a clear negative association with the probability of cross-species transmission (Fig. 4B, blue, p<0.001, coef. 95% CI (−7.633, −2.213)). Importantly, as already discussed, the strength of this association may be inflated by instances of lineage tracking (virus/host cospeciation). However, it is well established that increasing host genetic distance is associated with a higher species barrier; as previously documented in the literature (reviewed in [34]), we expect this is largely due to the divergence of host restriction factor genes, which are key components of the innate immune system. Functional assays of these host restriction factors against panels of SIVs, while outside the scope of this study, will be important for further identifying the molecular bases of the species barriers that have led to the transmission patterns identified here.

### Origins of HIV-1 and HIV-2

#### Epidemiological factors were key to the early spread of HIV

Understanding the underlying dynamics of lentiviral CST provides important biological context to the transmissions that generated the HIV pandemic. As discussed above, our results support a view of lentiviral cross-species transmission as a rare event. Notably, only two lentiviruses have crossed the high species barrier from Old World monkeys into hominids: SIV*smm* and the recombinant SIV*cpz*. Both HIV-1 and HIV-2 have arisen in human populations in the last century [5,35]. While it is possible that this has occurred by chance, even without increased primate exposure or other risk factors, we nevertheless find it striking that humans would acquire two exogenous viruses within such a short evolutionary timespan.

Examining this phenomenon more closely, the history of HIV-2 is enlightening. HIV-2 has been acquired through at least 8 *independent* spillover events from sooty mangabeys [5]. Notably, 6 of these transmissions have resulted in only a single observed infection (spillovers) [7,8,36]; only 2 of these events have established sustained transmission chains and successfully switched hosts to become endemic human pathogens [37–40]. Given this pattern, it is conceivable that there have been many isolated introductions of lentiviruses into humans over the past 200,000 years [14,41]. However, these other viral exposures did not result in new endemic human pathogens either because of species-specific immune barriers, non-conducive epidemiological conditions, or a combination thereof. The rapid and repeated emergence of HIV-1 and HIV-2 is on a timescale more congruent with changes in epidemiological conditions than mammalian evolution, perhaps emphasizing the importance of the concurrent changes in human population structure and urbanization in facilitating the early spread of the epidemic [35]. Significantly, though, this also highlights the importance of careful public health surveillance and interventions to prevent future epidemics of zoonotic viruses.

#### The exact origins of SIVcpz may not be identifiable

The ancestry of SIV*cpz* appears to be tripartite: the unknown ancestry of the 5’ end; the putatively SIV*rcm* or SIV*mnd-2* derived *int* and *vif* genes; and the putatively SIV*gsn/mon/mus* derived 3’ end of the genome. For any of these three portions of the genome, there are multiple evolutionary histories supported by available sequence data. However, it is possible that further sampling of lentivirus lineages (both known and currently undiscovered) will be able to definitively resolve the ancestry of SIV*cpz*. Alternatively, it may be that the ancestral virus(es) that gave rise to any of these three portions is extinct. In the case of the last two portions of the SIV*cpz* genome, it may be that the common ancestor of these putative genetic donors (SIV*rcm/mnd-2* and SIV*mon/mus/gsn*, respectively) was the true source. However, it is also a distinct possibility that SIV*cpz* has sufficiently diverged since its acquisition by chimpanzees such that its history is obscured.

#### Evolutionary time obscures the identity of the “original” primate lentivirus

Among primates, our results clearly illustrate that the vast majority of lentiviral lineages were originally acquired by cross-species transmission. It is intriguing to speculate as to which virus was the “original” source of all of these lineages. Because of its consistent position as the outgroup of primate lentiviral trees, there has long been a popular hypothesis in the field that SIV*col* was this original lentivirus among primates [27]. While SIV*col* is certainly the most evolutionary isolated extant lentivirus that has been sampled to date, this does not definitively place it as the ancestral lentivirus. Alternative scenarios (also noted by Compton *et al.*) include an extinct original lentiviral lineage (and/or primate host species) or an unsampled ancestral lentivirus. It is also plausible that another known extant lentivirus was the “original” lineage, but has diverged and/or recombined to such an extant that its origins are obscured.

#### Conclusions

Here, we have shown that lentiviruses have a far more extensive history of host switching than previously described. Our findings also demonstrate that the propensity of each lentiviral lineage to switch between distant hosts, or to spillover among related hosts, is highly variable. In examining specific lineages, our findings are consistent with the prevalent hypothesis that SIV*col* has evolved in isolation from other SIVs. Contrastingly, we have also demonstrated that the mosaic origins of SIV*cpz* are far more complex than previously recognized; the currently available sequence data is unable to resolve the ancestry of nearly half of the SIV*cpz* genome. Together, our analyses move closer to a full understanding of the pattern of cross-species transmission among primate lentiviruses, but additional efforts to obtain high quality viral genome sequences from under sampled lineages will be necessary to fully resolve the natural history of these viruses.

## Methods

All datasets, config files, documentation, and scripts used in this analysis or to generate figures are available publicly at <https://github.com/blab/siv-cst>.

### Datasets & Alignments

Lentiviral genomes are translated in multiple reading frames; we therefore utilized nucleotide sequence data for all analyses.

#### Recombination

For all recombination analyses (GARD, R^2^, rSPR), we used the 2015 Los Alamos National Lab (LANL) HIV/SIV Compendium [18]. The compendium is a carefully curated alignment of high-quality, representative sequences from each known SIV lineage and each group of HIV-1 and HIV-2. We reduced the overrepresentation of HIV in this dataset, but maintained at least one high quality sample from each group of HIV. In total, this dataset contains 64 sequences from 24 hosts (1-10 sequences per host). Maximum likelihood trees for each segment—used for the rSPR analysis and displayed in Figure 1—were built with RAxML v.8.2.9 [42] with the rapid bootstrapping algorithm under a GTR model with 25 discrete bins of site to site rate variation and were rooted to SIVcol.

#### Main & Supplemental Datasets

Primate lentiviral sequences were downloaded from the comprehensive LANL HIV sequence database [18]. Sequences from lineages known to be the result of artificial cross-species transmissions (SIVmac, -stm, -mne, and -wcm) were excluded. We also excluded any sequences shorter than 500 bases or that were flagged as problematic by standard LANL criteria. We grouped host subspecies together except for cases where there is a known specific relationship between host subspecies and virus lineage (chimpanzees and African green monkeys). To construct datasets with a more equitable distribution of sequences per host, we preferentially subsampled sequences from the LANL Compendium, followed by samples isolated from Africa (more likely to be primary sequences), and finally supplementing with samples isolated elsewhere (excluding experimentally generated sequences). For humans, mandrils and mustached monkeys, which are host to 2, 2 and 3 distinct viral lineages, respectively, we allowed a few additional sequences (if available) to represent the full breadth of documented lentiviral diversity. The “main” dataset consists of 24 host species, with 5-31 sequences per host (total N=422). As an alternative dataset to control for sampling bias and data availability, we also constructed a “supplemental” dataset with just 15 hosts but with more viral sequences per host (16 – 40 sequences per lineage, N=510).

#### Alignments

Alignments were done with the L-INSI algorithm implemented in mafft v7.294b [43]; the Compendium alignment was held fixed, with other sequences aligned to this template. Insertions relative to the fixed compendium were discarded. This alignment was then split along the breakpoints identified by GARD to yield the segment-specific alignments.

### Recombination

#### Topology-Based Analysis

Each portion of the genome was analyzed with the Genetic Algorithm for Recombination Detection (GARD), implemented in HyPhy v.2.2.0 [17], with a nucleotide model selected by HyPhy’s NucModelCompare package (#012234) and general discrete distribution (3 bins) of site variation.

Computational intensity was eased by analyzing the recombination dataset in 3kb long portions, with 1kb overlaps on either end (e.g., bases 1:3000, 2000:5000, 4000:7000, etc.). To control for sites’ proximity to the ends of the genomic portion being analyzed, this was repeated with the windows offset (e.g., bases 1:2500, 1500:4500, 3500:6500, etc.). In total, this resulted in every site of the alignment being analyzed at least two-fold, with at least one of these replicates providing 500-1500 bases of genomic context on either side (other than at the very ends of the total alignment). Disagreement between window-offset replicates for a given breakpoint were minimal and almost always agreed within a few bases, with two exceptions: offset replicates for the breakpoint between segments 7 and 8 disagreed by 529 bases, and offset replicates for the breakpoint between 8 and 9 disagreed by 263 bases. In these instances, we used the average site placement.

#### Sitewise Linkage Analysis

Biallelic sites were identified across the genome (ignoring gap characters and polymorphisms present at <5%). These biallelic sites were compared pairwise to generate an observed and expected distribution of haplotypes (combinations of alleles between the two sites), and assessed with the statistic R^2^=χ^2^/(d*N), where χ^2^ is chi-square, d is the degrees of freedom, and N is the number of haplotypes for this pair of sites [44]. This statistic follows the canonical interpretation of R^2^, i.e., if the allele at site 1 is known, how well does it predict the allele at site 2 (0 indicating no linkage between sites, and 1 indicating perfect linkage).

#### Segment Linkage Analysis

Segment linkage was assessed by comparing the similarity of tree topologies between segments. This was done with the pairwise method in the Rooted Subtree-Prune-and-Regraft (rSPR) package [45], which measures the number of steps required to transform one topology into another. Segment pairs with similar topologies have lower scores than segments with less similar topologies.

### Phylogenies & Discrete Trait Analysis

#### Empirical tree distributions

For each segment alignment, a posterior distribution of trees was generated with BEAST v. 1.8.2 under a general time reversible substitution model with gamma distributed rate variation and a strict molecular clock [46]. We used a Yule birth-death speciation model tree prior [47]; all other priors were defaults. Trees were estimated using a Markov chain Monte Carlo (MCMC). The main dataset was run for 25 million steps, sampled every 15,000 states after discarding the first 10% as burnin, resulting in ~1600 trees per segment in the posterior distribution. Convergence was determined by visually inspecting the trace in Tracer v. 1.6.0 [48]. All but two effective sample size (ESS) values (segment 2: AG substitution rate [ESS=164] and frequencies 4 [ESS=189]) were well over 200. The supplemental dataset was run for 65 million steps, sampled every 15,000 states after discarding the first 10% as burnin (~3900 trees per segment in the posterior distribution). Convergence was again determined in Tracer. All ESS values except for the AG substitution rate in segment 2 (ESS=190) were well above 200.

#### Discrete Trait Analysis

These tree distributions were used to estimate the transmission network. As in Faria *et al.* [22], we model hosts as states of a discrete trait, and model cross-species transmission as a stochastic diffusion process among *n* host states. We use a non-reversible state transition matrix with *n* × (*n−1*) individual transition parameters [49]. We also utilize standard methodology in using a Bayesian stochastic search variable selection process to encourage a sparse network, implemented as in [23]. An exponential distribution with mean=1.0 was used as a prior on each of the pairwise rate parameters (552 in total for the main dataset with 24 hosts; 210 parameters in total for the supplemental dataset with 15 hosts). The prior placed on the *sum* of the binary indicator variables corresponding to each pairwise rate parameter (i.e., the number of transitions that are non-zero) was an exponential distribution with mean=20. Convergence was again assessed by visually inspecting the log files in Tracer. The main dataset MCMC chain was run for 25 million steps and sampled every 5,000 steps after discarding the first 5 million steps as burnin. The supplemental dataset MCMC chain was run for 45 million steps and sampled every 5,000 steps after discarding the first 20 million steps as burnin. For both datasets, more than 97% of the rate, indicator, and actualRate (stepwise rate*indicator values) parameters had an ESS well over 200 (most > 1000).

Statistical support for each transmission is summarized as a Bayes factor (BF), calculated by comparing the posterior and prior odds that a given rate is non-zero (i.e., that there has been *any* transmission between a given pair of hosts) [23]. The ancestral tree likelihoods of each of the 12 tree distributions contribute equally to the inference of a shared transmission rate matrix. However, not every lineage has recombined along every breakpoint, which means that the tree likelihoods from each segment are not fully statistically independent. Thus, we report conservative estimates of BF by dividing all BF values by 12.

## Acknowledgements

We thank Michael Emerman, Janet Young, Gytis Dudas, Chris Whidden, and the Emerman and Bedford labs for their help. We also acknowledge the Los Alamos National Labs HIV database, which provided the curated data for this study. T.B. is a Pew Biomedical Scholar and his work is supported by NIH award R35 GM119774-01. S.B. is a NSF Graduate Research Fellow and her work is supported by NSF grant DGE-1256082 and NIH training grant 5-T32-CA-80416-17.

---

**Figure S1:**
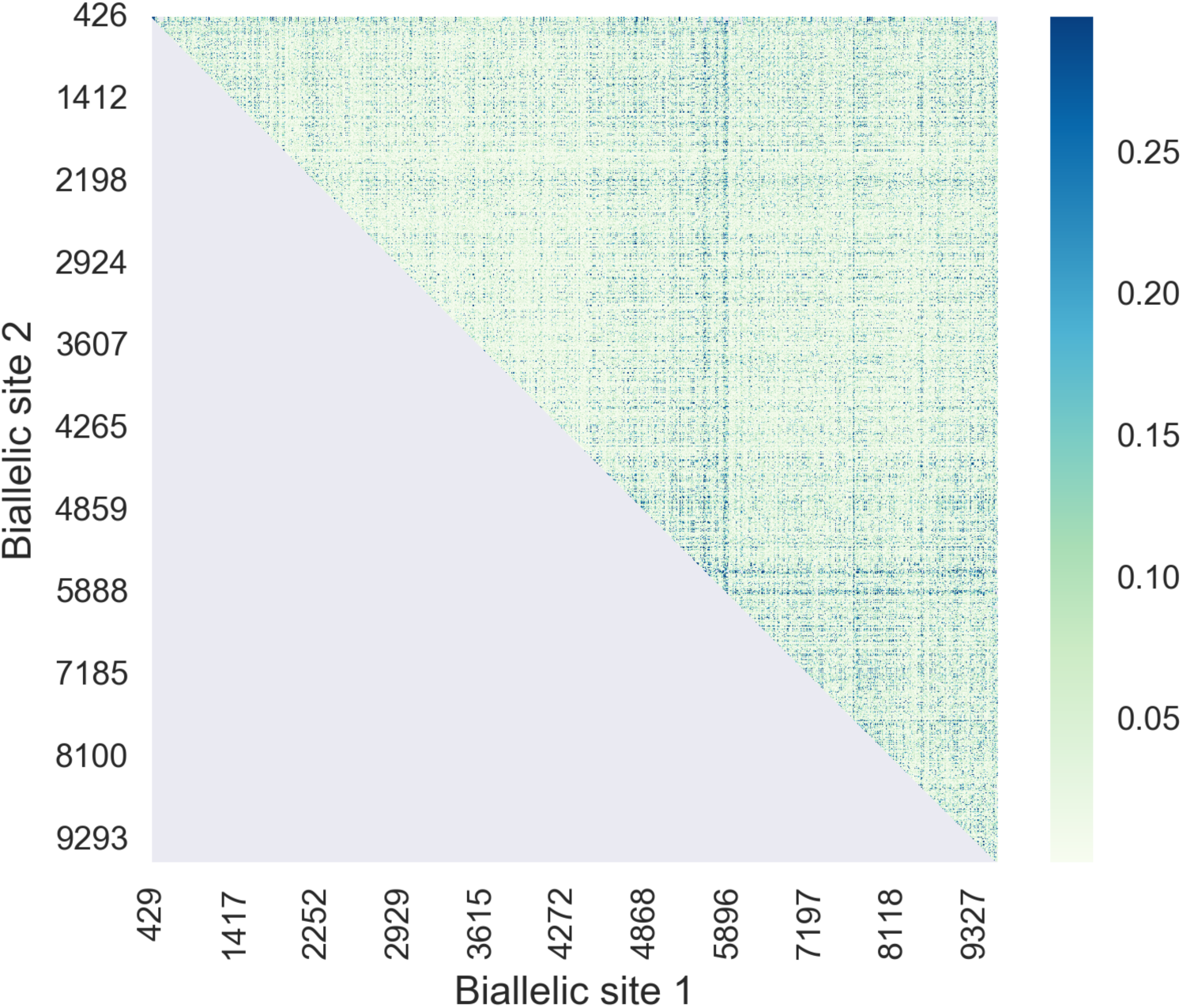
Extensive divergence makes sitewise measures of genetic linkage ineffective. For pairs of biallelic sites (ignoring rare variants), R^⋀^2 was used to estimate how strongly the allele in one site predicts the allele in the second site, with values of 0 indicating no linkage and 1 indicating perfect linkage. The mean value of R^⋀^2 was 0.044, indicating very low levels of linkage overall.

**Figure S2:**
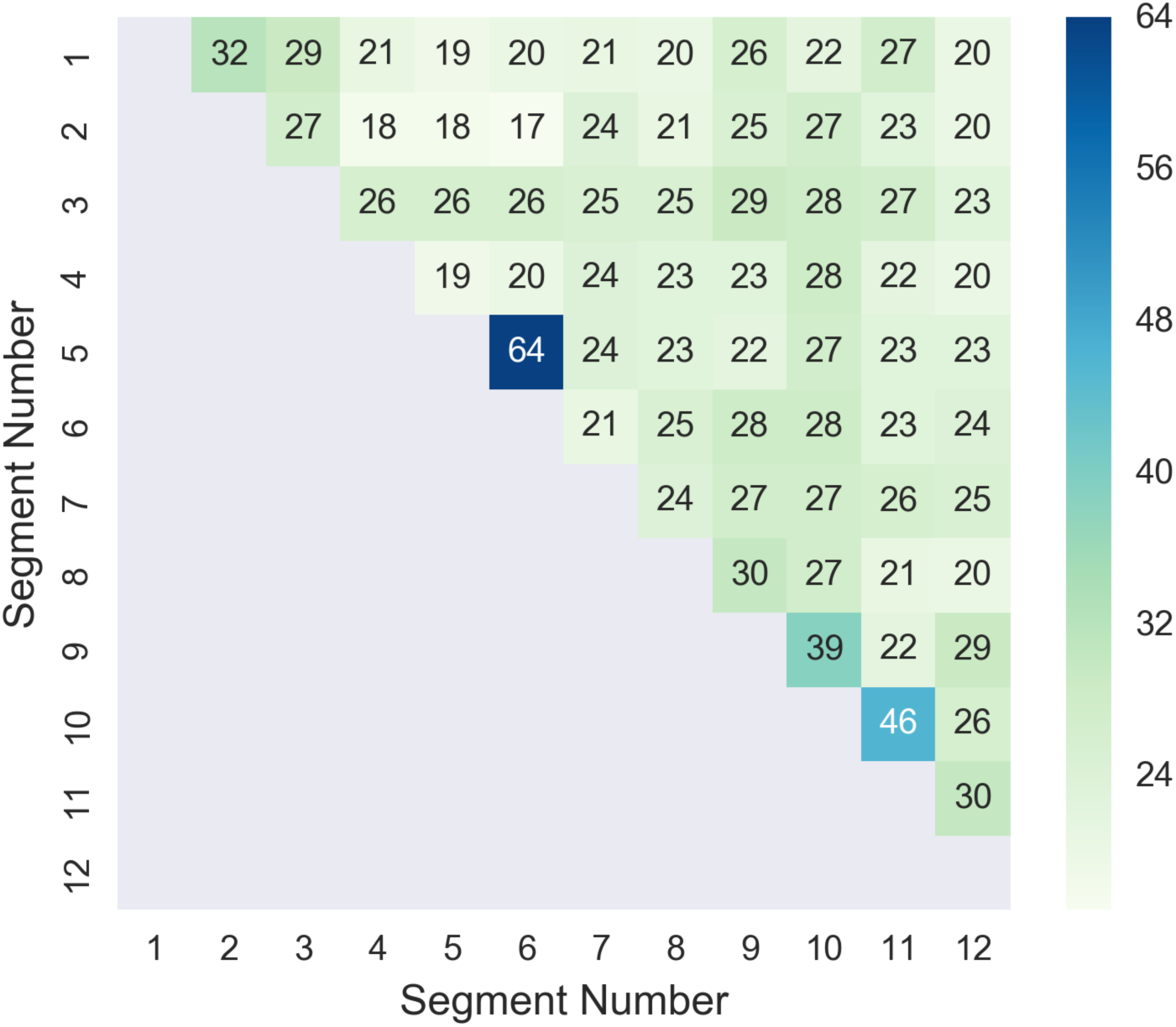
No evidence of linkage between nonadjacent segments of the SIV genome. The alignment used for GARD analyses (LANL compendium with HIV overrepresentation reduced) was split along the breakpoints identified by GARD to yield the 12 genomic segments, and a maximum likelihood tree was constructed for each. The number of steps required to turn one tree topology into another was assessed for each pair of trees with the Rooted Subtree-­‐Prune-­‐and-­‐Regraft (rSPR) package. Segment pairs with similar topologies have lower scores than segments with less similar topologies.

**Figure S3.**
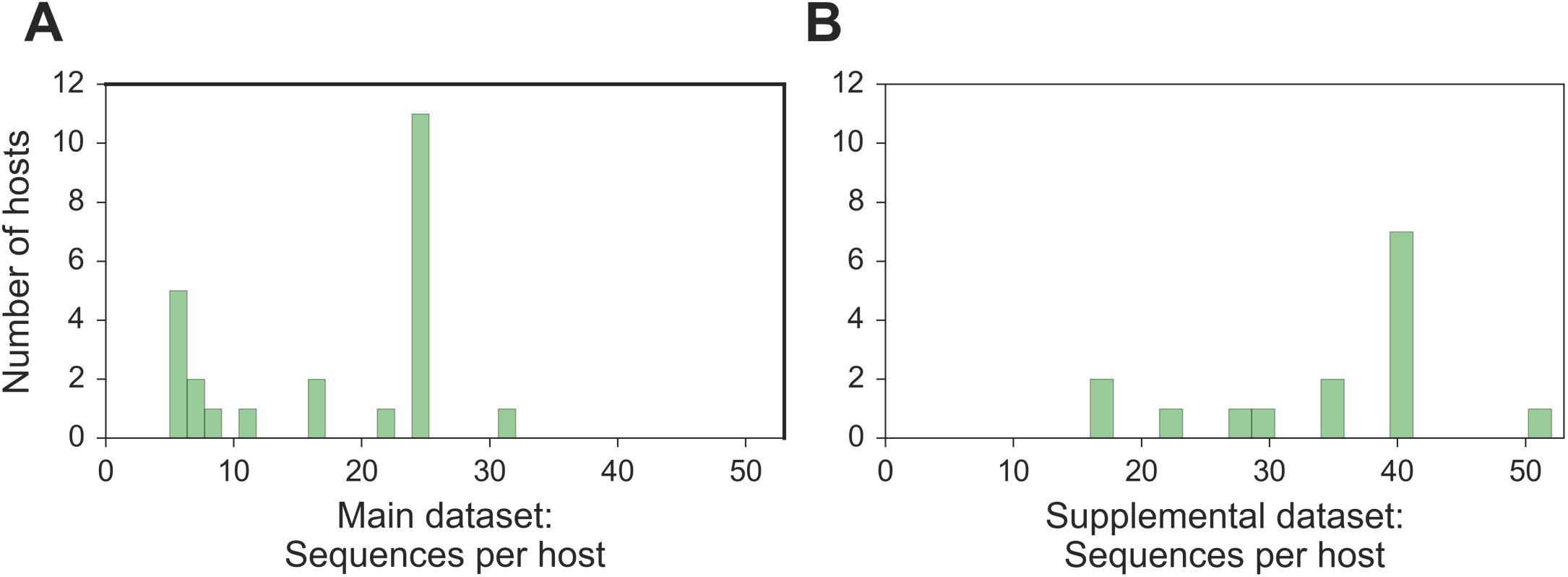
Distribution of the number of sequences per host included in analyses. A. All available high-­‐quality lentivirus sequences were randomly subsampled up to 25 sequences per host for the main dataset. We included the 24 hosts with at least 5 sequences available in this dataset. B: For the supplemental dataset, we randomly subsampled up to 40 sequences per host, and included the 15 hosts with at least 16 sequences available in this dataset. For both datasets, a small number of additional sequences were permitted for the few hosts that are infected by multiple viral lineages in order to represent the full breadth of known genetic diversity of lentiviruses in each host population.

**Figure S4:**
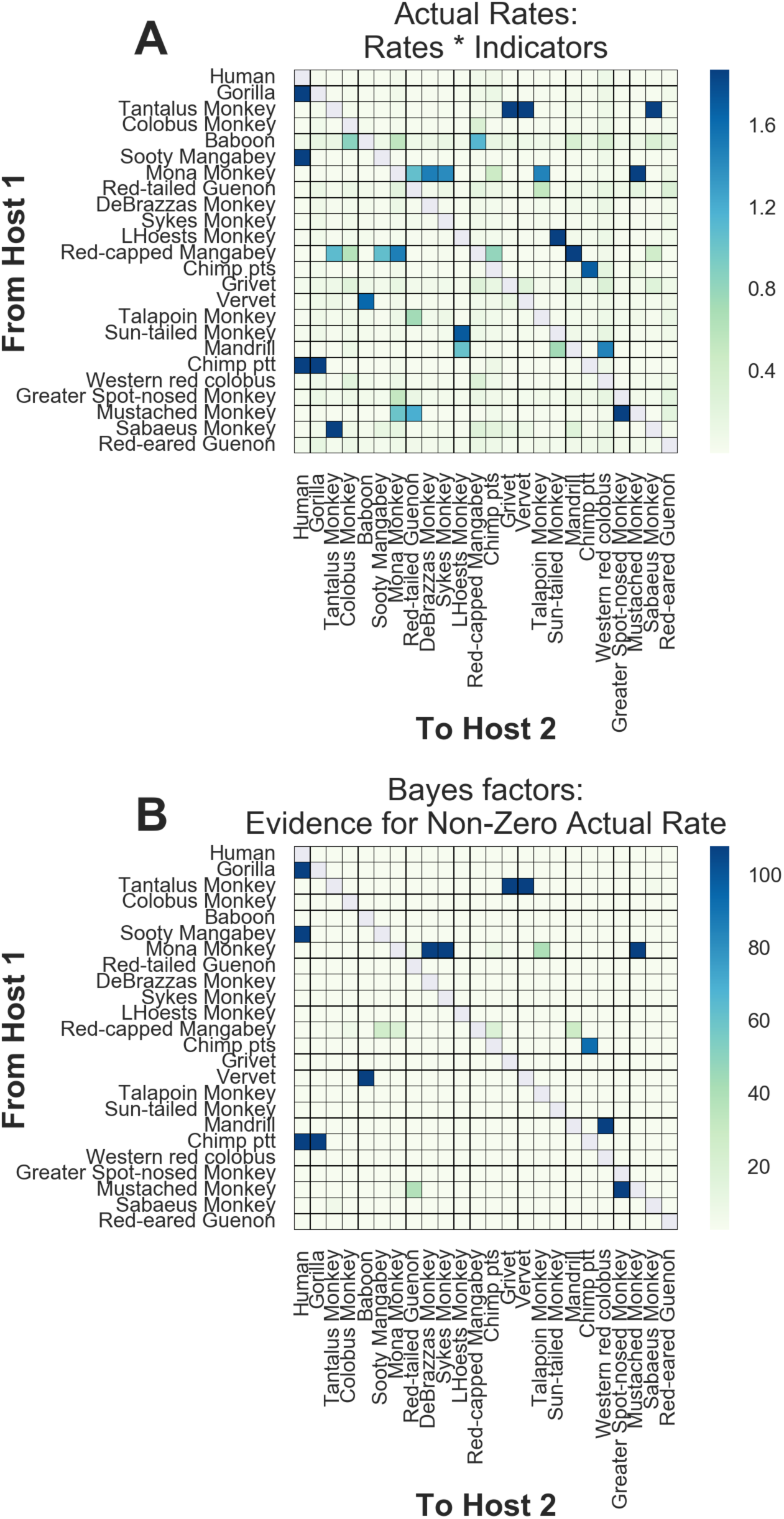
Actual rates and Bayes factors for main dataset discrete trait analyses. Values for the asymmetric transition rates between hosts, as estimated by the CTMC, were calculated as rate * indicator (element-­‐wise for each state logged). We report the average posterior values above. Bayes factors represent a ratio of the posterior odds / prior odds that a given actual rate is non-­‐zero. Because each of the 12 segments contributes to the likelihood, but they have not evolved independently, we divide all Bayes factors by 12 and report the adjusted values above (and throughout the text).

**Figure S5:**
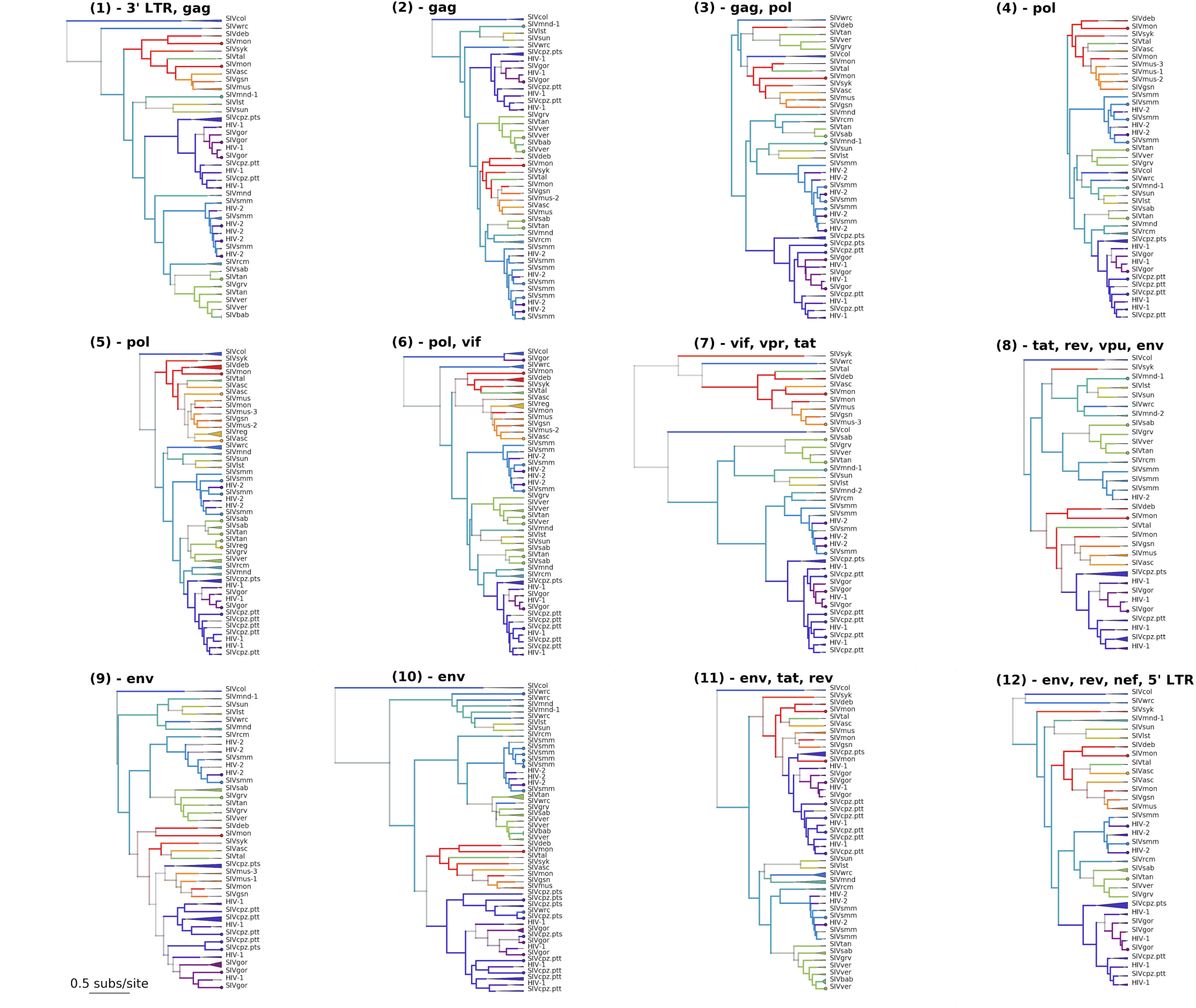
Maximum clade credibility trees for each of the 12 GARD-identified genomic segments of the lentiviral genome. Tips are color coded by known host state; branches and internal nodes are color coded by inferred host state, with color saturation indicating the confidence of these assignments. Monophyletic clades of viruses from the same lineage are collapsed, with the triangle width proportional to the number of represented sequences.

**Figure S6:**
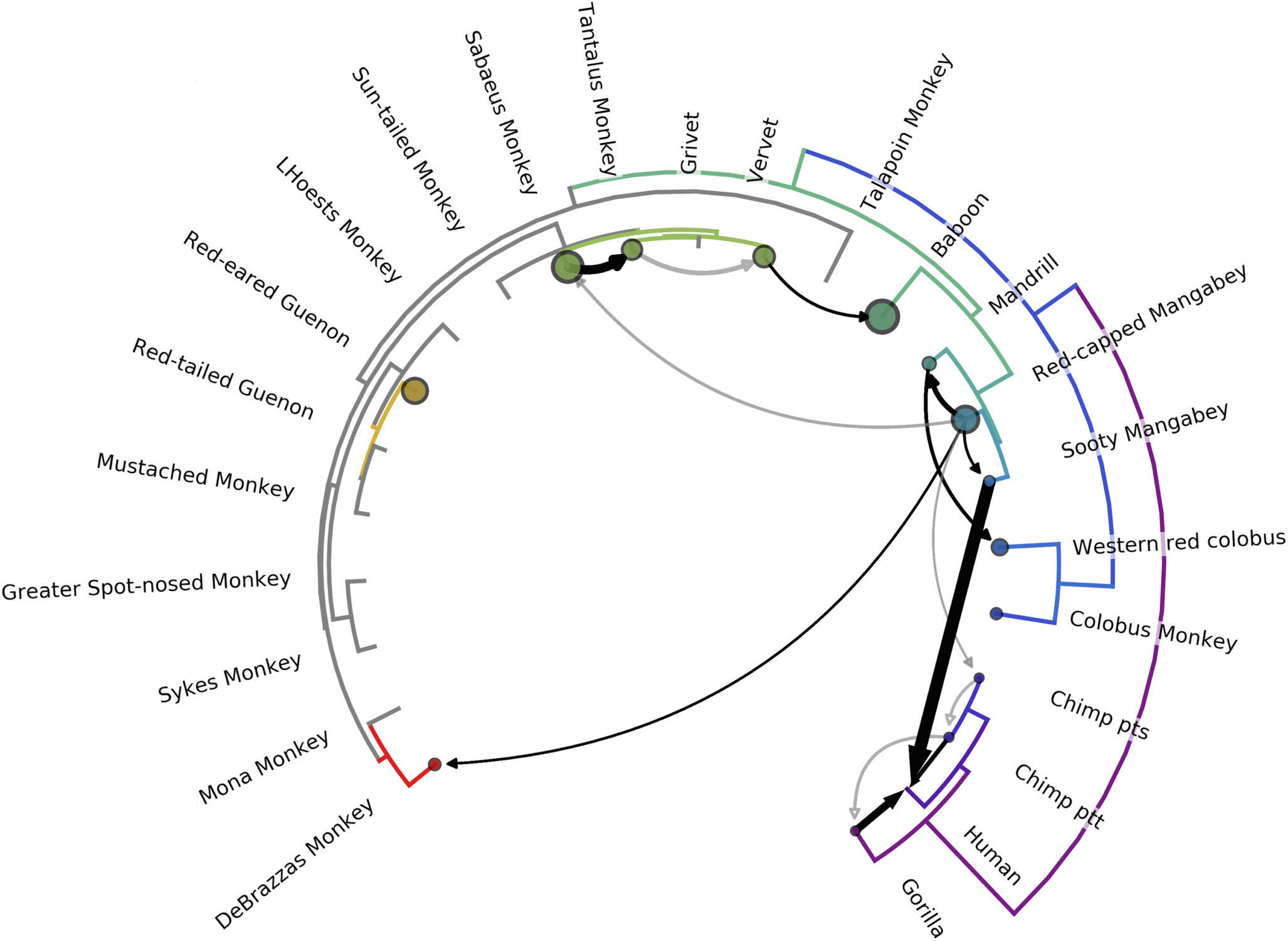
Most lentiviruses are the product of ancient cross-species transmissions (supplemental dataset). The phylogeny of the host species’ mitochondrial genomes forms the outer circle (gray: not included in supplemental dataset). Arrows with filled arrowheads represent transmission events inferred by the model with Bayes’ factor (BF) >= 3.0; black arrows have BF >= 10, with opacity of gray arrows scaled for BF between 3.0 and 10.0. Transmissions with 2.0 <= BF < 3.0 have open arrowheads. Width of the arrow indicates the rate of transmission (actual rates = rates * indicators). Circle sizes represent network centrality scores for each host. Transmissions from chimps to humans; chimps to gorillas; gorillas to humans; sooty mangabeys to humans; sabaeus to tantalus; and vervets to baboons have been previously documented. To our knowledge, all other transmissions illustrated are novel identifications.

**Figure S7:**
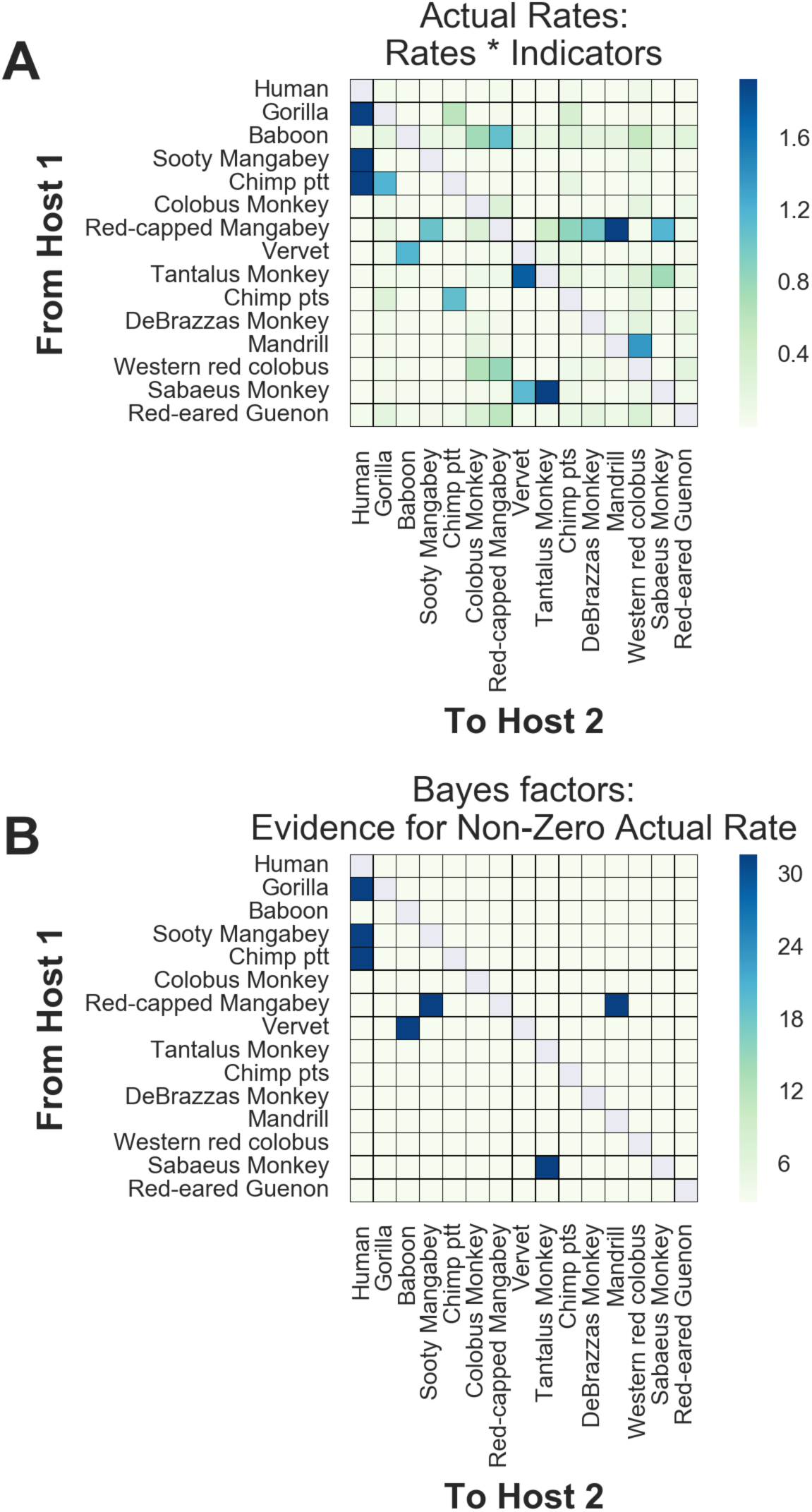
Actual rates and Bayes factors for supplemental dataset discrete trait analyses. Values for the asymmetric transition rates between hosts, as estimated by the CTMC, were calculated as rate * indicator (element-­‐wise for each state logged). We report the average posterior values above. Bayes factors represent a ratio of the posterior odds / prior odds that a given actual rate is non-­‐zero. Because each of the 12 segments contributes to the likelihood, but they have not evolved independently, we divide all Bayes factors by 12 and report the adjusted values above (and throughout the text).

**Figure S8:**
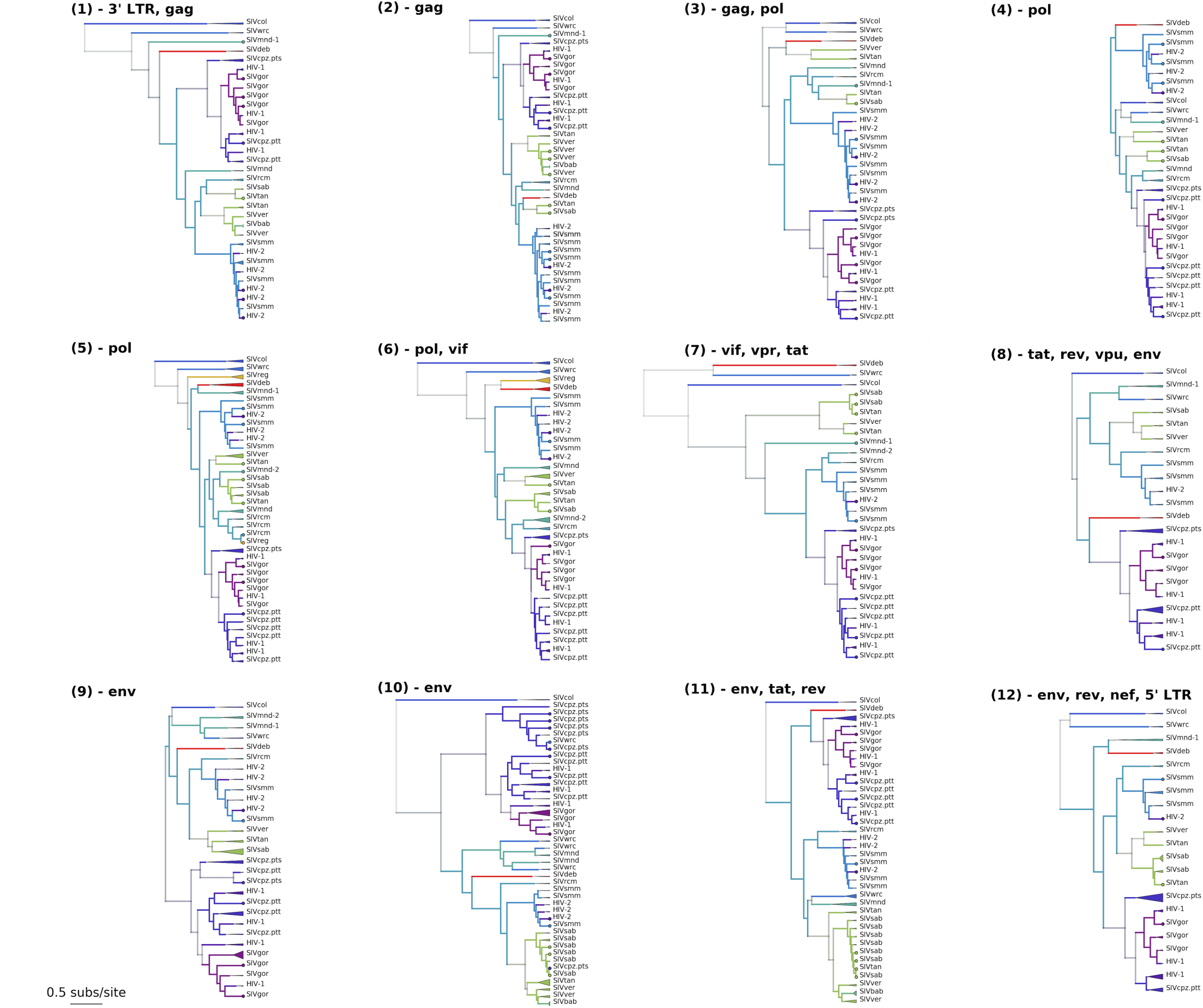
Maximum clade credibility trees for each of the 12 GARD-­‐identified genomic segments of the lentiviral genome (supplemental dataset) Tips are color coded by known host state; branches and internal nodes are color coded by inferred host state, with color saturation indicating the confidence of these assignments. Monophyletic clades of viruses from the same lineage are collapsed, with the triangle width proportional to the number of represented sequences.

**Figure S9:**
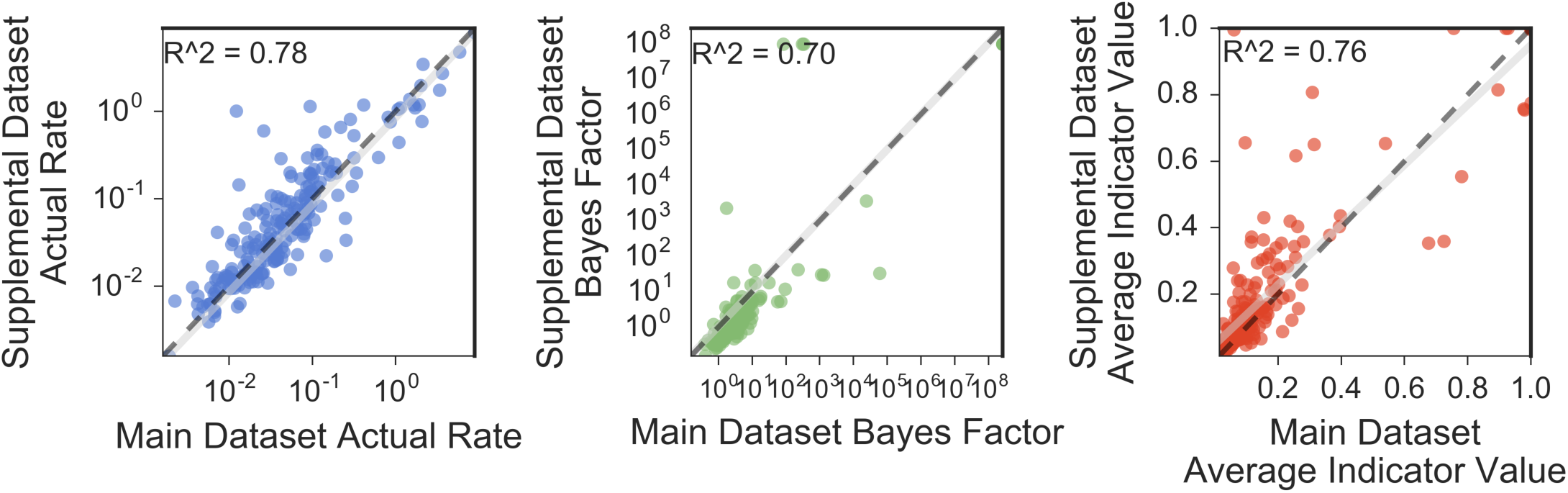
Comparison of Main and Supplemental Dataset Discrete Trait Analysis Results. Each datapoint represents on of the 210 possible transmissions between each pair of the 15 hosts present in both datasets. The black dashed line shows y=x; the linear regression and 95% CI are shown in gray.

